# A heterocyclic compound inhibits viral release by inducing cell surface BST2/Tetherin/CD317/HM1.24

**DOI:** 10.1101/2024.05.03.592399

**Authors:** Perpetual Nyame, Akihiro Togami, Tomofumi Yoshida, Takuya Masunaga, MST Monira Begum, Hiromi Terasawa, Nami Monde, Yurika Tahara, Reiko Tanaka, Yuestu Tanaka, Joyce Appiah-Kubi, Wright Ofotsu Amesimeku, Md Jakir Hossain, Masami Otsuka, Kazuhisa Yoshimura, Terumasa Ikeda, Tomohiro Sawa, Yorifumi Satou, Mikako Fujita, Yosuke Maeda, Hiroshi Tateishi, Kazuaki Monde

**Author notes:** Correspondence: Kazuaki Monde, Department of Microbiology, Faculty of Life Sciences, Kumamoto University, Kumamoto 860-8556, Japan E-mail address Hiroshi Tateishi, Medicinal and Biological Chemistry Science Farm Joint Research Laboratory, Faculty of Life Sciences, Kumamoto University, Kumamoto 860-8556, Japan.

## Abstract

The introduction of combined antiretroviral therapy (cART) has greatly improved the quality of life of human immunodeficiency virus type 1 (HIV-1)-infected individuals. Nonetheless, the ever-present desire to seek out a full remedy for HIV-1 infections makes the discovery of novel antiviral medication compelling. Owing to this, a new late-stage inhibitor, Lenacapavir/Sunlenca, an HIV multi-phase suppressor, was clinically authorized in 2022. Besides unveiling cutting-edge antivirals inhibiting late-stage proteins or processes, newer therapeutics targeting host restriction factors hold promise for the curative care of HIV-1 infections. Notwithstanding, bone marrow stromal antigen 2 (BST2)/Tetherin/CD317/HM1.24, which entraps progeny virions is an appealing HIV-1 therapeutic candidate. In this study, a novel drug screening system was established, using the Jurkat/Vpr-HiBiT T cells, to identify drugs that could obstruct HIV-1 release; the candidate compounds were selected from the Ono Pharmaceutical compound library. Jurkat T cells expressing Vpr-HiBiT were infected with NL4-3, and the amount of virus release was quantified indirectly by the amount of Vpr-HiBiT incorporated into the progeny virions. Subsequently, the candidate compounds that suppressed viral release were used to synthesize the heterocyclic compound, HT-7, which reduces HIV-1 release with less cellular toxicity. Notably, HT-7 increased cell surface BST2 coupled with HIV-1 release reduction in Jurkat cells but not Jurkat/KO-BST2 cells. Seemingly, HT-7 impeded simian immunodeficiency virus (SIV) and severe acute respiratory syndrome coronavirus 2 (SARS-CoV-2) release. Concisely, these results suggest that the reduction in viral release, following HT-7 treatment, resulted from the modulation of cell surface expression of BST2 by HT-7.

**Importance:** A collection of scientific strategies has been revealed to find long-term cure for HIV-1 infection. One of these techniques, the therapeutic approach, involves harnessing late events that are not targeted by current medication. The regulator of HIV-1 assembly and release, the HIV-1 Gag protein, has emerged as a prospective inhibitor. We set up a high-efficiency, economically viable, and facile screening system for the identification of late-stage inhibitors. Herein, we discovered a heterocyclic compound that inhibits HIV-1 release. This newly high-performance testing technique can be employed in virological research for investigating HIV- 1 late-stage processes.

## Introduction

Despite enormous scientific efforts to eradicate the human immunodeficiency virus type 1 (HIV-1), HIV-1 is still considered a worldwide disease burden [1]. Undoubtedly, the commencement of combined antiretroviral therapy (cART) in 1996 tremendously improved the living condition of HIV-1-infected individuals by repressing viral replication, and thereby attaining undetectable viremia [2]. To date, the Food and Drug Administration (FDA) has clinically approved approximately 43 HIV-1 antiretroviral drugs [3]. However, until recently, protease inhibitors were the only existing cARTs to target the late stage of the viral life cycle.

While the early phase of the viral life cycle commences with the attachment of the virus to the susceptible host cells and lasts up to the integration of viral DNA into the host genome, the late stage encompasses gene expression, viral release, and maturation of nascent virions. About ten protease inhibitors (e.g., darunavir) [4, 5] and the currently approved long-acting capsid inhibitor (lenacapavir) [6] hinder the formation of mature particles. So far, several compounds targeting virus release have been discovered [7–11]. Notable among these compounds are those that occlude Gag membrane binding (compound 7) [7], assembly of Gag (capsid assembly inhibitors; CAI peptide) [8], and viral pinch-off (cyclin peptide 11) [9].

Nevertheless, these compounds have not been approved for clinical use. Concurrently, the demand to enhance cART options [6] has made it necessary to develop drugs that act in the late stage of the viral life cycle.

HIV-1 Gag consists of four major domains: matrix (MA), capsid (CA), nucleocapsid (NC), and late domain (p6), and is predominantly associated with the release and maturation of the progeny virus at the late stage [12, 13]. The MA facilitates the targeting and binding of Gag to phosphatidylinositol 4,5-bisphosphate [14, 15] through the exposure of the myristate group [16–19]. Following membrane binding, MA engages the enveloped glycoprotein in the growing virions [20, 21]. While CA is crucial in the formation of an immature Gag lattice through Gag multimerization [22, 23], the NC packages viral genomic RNA into the virions [24]. Not only does the p6 attain viral pinch-off through the cellular endosomal sorting complex required for transport machinery, but it also packages the viral accessory protein (Vpr) into the virions [25, 26]. Given the role of HIV-1 Gag in the late stage of the viral life cycle, any compound that targets its functioning would undoubtedly be a major inhibitor of this phase of the life cycle.

Lately, there has been a profound interest in exploiting cellular restriction factors in the design and development of new antiviral drugs [27, 28]. The most studied cellular antiviral regulator against HIV-1 replication is BST2 (Bone marrow stromal cell antigen 2)/Tetherin/CD317/HM1.24. BST2 is an interferon-inducible protein expressed by numerous cell types that exhibit its inhibitory effects on viral release [29, 30]. With the help of its components sited in the plasma membrane, BST2 confines nascent virions on the cell surface, thereby accomplishing its release inhibitory effect on HIV-1 replication [31]. The extraordinary BST2’s structural component (transmembrane, coiled-coil, and GPI anchor domain) enables it to entangle budding virions on the cell membrane [29, 32]. Earlier studies illustrated that BST2 colocalizes with HIV-1 Gag on the cell surfaces as well as in endosomes, thus showcasing BST2’s release inhibitory impact on the cell surface and within cells [29]. On the other hand, viral accessory proteins have perpetually evolved interfering mechanisms against host antiviral factors, invariably accelerating their replication in susceptible hosts [33]. Regarding BST2, diverse viral proteins nullify its influence on viral replication. For instance, viral Vpu and Env equilibrate BST2’s impact on HIV-1 and -2 replication. Conversely, the simian immunodeficiency virus (SIVcpz, SIVtan, and other SIV) neutralizes BST2 activity by Vpu, Env, and Nef proteins [30, 34]. The ability of BST2 to combat enveloped viruses and its species specificity makes it a plausible candidate for HIV-1 therapy. Unquestionably, a compound with the potency of modulating BST2 may restrain HIV-1 release. To date, a compound aimed at harnessing BST2 activities against HIV-1 has not yet received clinical approval.

Here, we established a novel system using the Jurkat/Vpr-HiBiT T cells to screen for drugs that hinder virus release. The procedure and cost of this screening system are not laborious and costly compared to those of previous systems, such as the p24 enzyme-linked immunosorbent assay (ELISA). Using the screening system, candidate compounds obtained from the Ono Pharmaceutical compound library that restrain HIV-1 release were selected. A derivative compound HT-7 attenuated viral pinch-off by triggering the production of cell surface BST2 with negligible cytotoxic effect. In a nutshell, this study unveils a high-performance technique for exploring HIV-1 late-stage inhibitors. This technique revealed HT-7 to enhance the T-cell surface expression of BST2, which interferes with the efficiency of viral release.

## Materials and Methods

### Plasmids

The HIV-1 molecular clone pNL4-3/KFS contains a frameshift mutation that disrupts Env expression [35]. pNL4-3/GagVenus, encoding Gag fused to the mVenus variant of yellow fluorescent protein (YFP), fusion at the C terminus of Gag, has been described previously [17]. The pNL4-3/Fyn(10)fullMA/GagVenus and pNL4- 3/Fyn(10)6A2T/GagVenus were elucidated in earlier studies [17, 36] (a kind gift from A. Ono). The Vpr protein was tagged with a 11 amino acid peptide tag (HiBiT) using PCR amplification. The sequences for the first and second reverse primers were respectively CGGCTGGCGGCTGTTCAAGAAGATTAGCTAGGTCGACCAGCTGTG and GAAATGGAGCCAGTAGATCCGTGAGCGGCTGGCGGCTGTTCAAGA. The first reverse primer encodes the HiBiT tag and SalI restriction site (SalI). The second reverse primer also encodes the HiBiT tag in addition to HIV-1 Vpr. The PCR product (Vpr-HiBiT) was then inserted into pRDI292 using the restriction enzymes BamHI and SalI. The U3 region was inserted into the 5′ UTR of pRDI292 to reconstruct the intact 5′LTR, thus pRDI292/Vpr-HiBiT encodes the LTR-driven Vpr-HiBiT and SV40-driven puromycin-resistance genes. To avoid packaging the pRD292/Vpr-HiBiT genes into the HIV-1 virions, the packaging signal was removed from the pRDI292/Vpr-HiBiT construct.

### Cells

293T cells were cultured in Dulbecco’s modified Eagle’s medium (DMEM; Sigma-Aldrich) supplemented with 10% fetal bovine serum (FBS) (NICHIREI). While Jurkat/Vpr-HiBiT cells were cultured in RPMI1640 medium (Gibco) with 10% FBS and 1 μg/ml puromycin (InvivoGen) (RPMI-10), VeroE6/TMPRSS2 cells (VeroE6 cells stably expressing human TMPRSS2; JCRB Cell Bank, JCRB1819) [37] were maintained in DMEM (low glucose, Wako) containing 10% FBS, G418 (1 mg/ml; Wako), and 1% penicillin/streptomycin (Wako) (DMEM-10).

### Generation of Jurkat/Vpr-HiBiT and 293T/Vpr-HiBiT cell lines

pRDI292/Vpr-HiBiT was digested and linearized using the restriction enzyme HpaI. The linearized plasmid was transfected into Jurkat T cells using the AMAXA Nucleofector (Lonza, Switzerland), with strict adherence to the manufacturer’s instructions. Similarly, following the manufacturer’s instructions, the linearized pRDI292/Vpr-HiBiT was transfected into 293T cells using Lipofectamine 3000 (Thermo Fisher Scientific). As described above, pRDI292/Vpr-HiBiT transduction using a lentiviral vector was unavailable since the packaging signal was removed. The transfected cells were cultured with puromycin for approximately 30 days, and the cells expressing the puromycin-resistance gene permanently were selected.

### Measurement of NanoLuciferase luminescence

Jurkat/Vpr-HiBiT cells were infected with VSV-G-pseudotyped NL4-3/KFS, whereas the 293T/Vpr-HiBiT cells were transfected with NL4-3/KFS. At 15 h post-incubation, the infected Jurkat/Vpr-HiBiT cells, and the transfected 293T/Vpr-HiBiT cells were similarly treated with the compounds/derivative compounds and incubated for two days. After two days of incubation, the supernatant from the infected Jurkat/Vpr-HiBiT and transfected 293T/Vpr-HiBiT cells was harvested. Subsequently, luminescence activity was analyzed by the Nano-Glo® HiBiT Lytic Detection System (Promega). Based on the manufacturer’s instructions, the cell supernatants were lysed with Nano-Glo® HiBiT Lytic Buffer. After adding LgBiT, the substrate was added to the lysed supernatants. The Luciferase activity was then measured using a GloMax® Discover System (GM3000).

### p24 Gag ELISA

The supernatant obtained from the NL4-3-infected Jurkat/Vpr-HiBiT cells and the pNL4-3-transfected 293T/Vpr-HiBiT cells was lysed with a lysis agent in p24 ELISA (ZeptoMetrix). Following the manufacturer’s instructions, the amount of Gag proteins in the supernatant was quantified using an in-house laboratory-manufactured kit described earlier [38] and a p24 ELISA kit (ZeptoMetrix).

### p27 Gag ELISA

A lysis agent in p27 ELISA (ZeptoMetrix) was employed to lyse the cell supernatant harvested from the SIVmac239-infected Jurkat/Vpr-HiBiT cells (SIVmac239; AIDS Research and Reference Reagent Program). The amount of Gag proteins (p27) in the supernatant was measured (ZeptoMetrix).

### Confocal microscopy

Jurkat/Vpr-HiBiT cells were infected with VSV-G-pseudotyped NL4-3/GagVenus. At 15 h post-infection, the compounds were added to the infected cells and incubated for one day. Later, the cells were fixed with 4% paraformaldehyde (PFA) (Wako) for 30 min at 4 °C and washed once with phosphate-buffered saline (PBS). Afterward, the cells were mixed with Fluoromount-G (Dako) and plated on the microscope slide (Matsunami). Images of 50 fields were recorded using a Zeiss LSM 700 laser-scanning confocal microscope.

### Flow cytometry analysis

VSV-G-pseudotyped NL4-3/GagVenus were harvested from the transfected 293T cells with pNL4-3 GagVenus, the GagPol expression vector pCMVNLGagPolRRE, and the VSV-G expression vector pHCMV-G, as reported previously [39]. Jurkat/Vpr-HiBiT cells were infected with the VSV-G-pseudotyped NL4-3/GagVenus. At 15 h post-infection, the infected cells were cultured with the compounds and incubated for 48 h. Approximately, 48 h post-infection, cells were fixed with 4% PFA for 1 h, washed twice with PBS, and then blocked for 1 h with 2% bovine serum albumin (BSA). Later, the blocked cells were washed twice with PBS and then stained with anti-mouse PSGL-1 (KPL1; Santa Cruz Biotechnology) and anti-mouse BST2 antibody (E-4; Santa Cruz Biotechnology) in a ratio of 1:100. After 1 h of incubation at 4°C, cells were washed three times with PBS and then stained with anti-mouse Alexa Fluor 546 antibody (Invitrogen) in a proportion of antibody: 2% BSA of 1:100. An hour post-infection, stained cells were washed twice and suspended in PBS. The fluorescence signal (YFP) was analyzed using flow cytometry (FACSCalibur, BD Biosciences).

### Establishment of BST2 knockout cells

A gRNA oligonucleotide (CGATTCTCACGCTTAAGACC) annealing to exon 4 of the ORF of the human BST2 gene was designed and cloned into lentiCRISPR v2 (Addgene). Next, the ribonucleoprotein complex, composed of the designed gRNAs and lentiCRISPR v2 was transfected into 293T cells. After 2 days of transfection, cell supernatants were pooled, filtered, and centrifuged at 13200 × *g* for 1 h at 4 °C. Viral pellets were resuspended in RPMI-10, followed by aspiration of the cell supernatant. Jurkat/Vpr-HiBiT cells were then transduced with the harvested virus. In parallel, 2 ml fresh pre-warmed DMEM-10 was added to the transfected 293T cells and incubated for two days. Afterward, the BST2-deficient (Jurakt/Vpr-HiBiT/KO-BST2) cells were selected upon treatment with 1 µg/ml puromycin.

Then, a day after initial drug treatment, 2 ml fresh pre-warmed RPMI-10 was added to the cells and re-treated with 1 µg/ml puromycin. At 48 h incubation, limiting dilutions were performed to achieve a monoclonal expansion of the BST2-deficient cell population. A preliminary selection of the Jurkat/Vpr-HiBiT/KO-BST2 cell clones by flow cytometry (FACSCalibur, BD Biosciences) assay was undertaken. Conclusively, the sequence information of BST2 ORF exon 4 in Jurkat/Vpr-HiBiT/KO-BST2 cells was analyzed by the Sanger sequencing technique.

### Sequencing analysis of extracted DNA

Genomic DNA was extracted from Jurkat/Vpr-HiBiT/KO-BST2 cells using a QIAGEN kit in compliance with the manufacturer’s protocol. The extracted DNA was amplified by Nested PCR using (Forward-CACAAAAGGATAACTTAGCC and Reverse- CCCCGCCCCCCTTCCCCAGC) as an outer primer pair and (Forward- CTTGGATTGGGGCGGTGCGG and Reverse-CACTGACCAGCTTCCTGGGA) as the inner primer pair. Hereinafter, PCR amplicons were purified in conformance with the QIAGEN PCR Purification Kit protocol. The purified PCR products were then ligated into pCR®-Blunt II-TOPO® vector (Invitrogen) and transformed into *E. coli* DH-5α competent cells (Takara). Ultimately, the sequence of the resulting recombinant plasmid was ascertained by Sanger sequencing techniques (Applied Biosystems™ 3130 Genetic Analyser), and the resulting data were decoded by ApE software (v2.0.36) created by M. Wayne Davis.

### SARS-CoV-2 D614G-bearing B.1.1 variant preparation

Severe acute respiratory syndrome coronavirus 2 (SARS-CoV-2) D614G-bearing B.1.1 variant (strain TKYT41838, GenBank accession no. LC606020) was propagated as previously described [40, 41]. Briefly, VeroE6/TMPRSS2 cells (5 × 10^6^ cells) were seeded in a T-75 flask the day before infection. From then on, the virus was diluted in virus dilution buffer [1 M HEPES, DMEM (low glucose), non-essential amino acid (Gibco), 1% penicillin/streptomycin], and the dilution buffer containing the virus was added to the flask after removing the initial medium. After 1 h of incubation at 37°C, the supernatant was replaced with 15 ml of 2% FBS/DMEM (low glucose), and further culture at 37°C was conducted until visible cytopathic effect (CPE) was observed. Upon CPE observation, the cell culture supernatant was collected, centrifuged at 300 × *g* for 10 min, and frozen at –80°C as a working virus stock. The titer of the prepared working virus was determined as the 50% tissue culture infectious dose (TCID_50_) [42, 43]. The day before infection, VeroE6/TMPRSS2 cells (10,000 cells) were seeded in a 96-well plate and infected with serially diluted working virus stocks. The infected cells were incubated at 37°C for 4 days and the appearance of CPEs in the infected cells was observed by a microscope. The value of TCID_50_/ml was calculated by the Reed-Muench method [44].

### SARS-CoV-2 B.1.1 infection

The day before infection, 1×10^4^ VeroE6/TMPRSS2 cells were plated in 96 well plates. The cells were inoculated with the SARS-CoV-2 B.1.1 variant (100 TCID_50_) and incubated at 37°C for 1 h. Thereafter, the supernatant was removed, cells were washed twice with fresh culture medium, and 200 µl of fresh culture medium containing HT-7 (final concentration 50 µM) was added and incubated at 37°C. Next, 15 μl of cell culture supernatant was harvested at the indicated time points (0, 24, 48, and 72 h, respectively), and the viral RNA copy number was quantified by RT-qPCR.

### RT-qPCR

RT-qPCR was performed as previously described [45–48]. Briefly, 5 μl of culture supernatant was mixed with 5 μl of 2 × RNA lysis buffer [2% Triton X-100 (Nacalai Tesque), 50 mM KCl, 100 mM Tris-HCl (pH 7.4), 40% glycerol, 0.8 U/μl recombinant RNase inhibitor (Takara)] and incubated at room temperature for 10 min. Next, 90 μl of RNase-free water was added and then 2.5 μl of diluted sample was used for real-time RT-PCR according to the manufacturer’s protocol with One step TB green PrimeScript PLUS RT-PCR Kit (Takara) and primers for the *nucleocapsid (N)* gene; Forward *N*, 5′-AGC CTC TTC TCG TTC CTC ATC-3′ and Reverse *N*, 5′-CCG CCA TTG CCA GCC ATT C-3′. The viral RNA copy number was standardized using a SARS-CoV-2 direct detection RT-qPCR kit (Takara). Fluorescent signals from resulting PCR products were acquired using a Thermal Cycler Dice Real Time System III (Takara).

### Statistical analysis

GraphPad Prism9 was the program used to perform statistical analysis on all of the data collected in this study. Besides one-way ANOVA with Sidak’s multiple comparison tests, comparison between data was also assessed by employing the two-way ANOVA with Tukey’s multiple comparisons. Likewise, unpaired t-tests and Pearson’s correlation coefficient were utilized in this study. The error bars indicate the mean of the standard deviation of repetitive independent experiments.

## Results

### Jurkat/Vpr-HiBiT is an established cell line for drug screening

To measure the amount of viral progeny, we established several Jurkat T cell lines expressing the reporter genes driven by HIV-1 Tat (Fig. 1A). In the nanoluciferase (NLuc; 19kDa)-expressing T cells, NLuc was secreted in the virus-free supernatant but not assembled into virions. Hence, Vpr-fused NLuc and Vpr-fused repetitive NLuc (2NLuc, 3NLuc, and 4NLuc) were transduced into Jurkat cells to accelerate the assembly of NLuc into the virions. However, the amount of all repetitive NLucs in virus-free supernatant was almost the same as that of NLuc in the cell supernatant. The Vpr fused with smaller peptide HiBiT (Vpr-HiBiT) in the virus-free supernatant was markedly reduced compared with that of Vpr-HiBiT in the cell supernatant. Consistently, the amount of p24 Gag assembled into virions was the same as that into the cell supernatant (Fig. 1B). The efficiency of the Vpr-HiBiT and p24 Gag assembly into the virions produced from 293T/Vpr-HiBiT cells was significantly lower than that in the cell supernatants (Figs. 1C and 1D). It indicates that approximately 25% of Vpr-HiBiT and p24 Gag secreted into the cell supernatant were not assembled into the virions, suggesting that Jurkat cells would be proper for drug screening. Firefly and renilla luciferase, which are large proteins (61kDa and 36kDa, respectively), were not assembled as HiBiT, suggesting that the smaller size of the reporter was key to the Vpr assembly into the virions (Fig. 1E). The amount of Vpr-HiBiT in the cell supernatant was evaluated by treatment with reverse transcriptase inhibitors (RT inhibitors: efavirenz and nevirapine) [49, 50] at pre-infection or 15 h post-infection using Jurkat/Vpr-HiBiT cells (Fig. 1F). As expected, the Vpr-HiBiT in the cell supernatant was markedly reduced by the treatment with RT inhibitors at pre-infection but not at 15 h post-infection. This result indicates that reverse transcription had been accomplished by 15 h post-infection. It suggests that the 15 h post-infection period was suitable for screening drugs targeting the late phase of the viral life cycle.

**Fig. 1.**
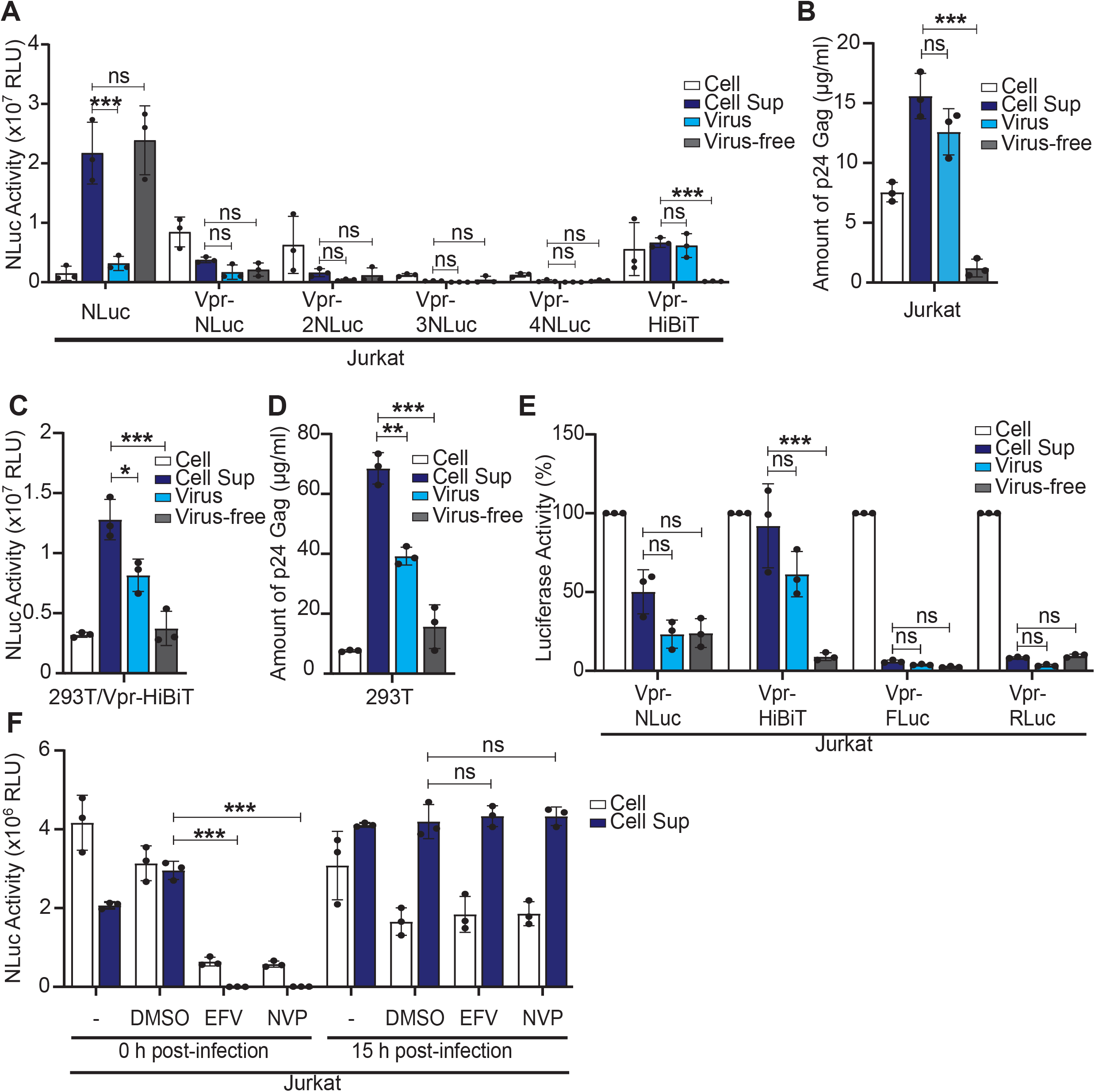
Vpr-HiBiT co-assembled with Gag into the virions. (A) Jurkat cells expressing the indicated nanoluciferase (NLuc), Vpr-fused NLuc, Vpr-fused repetitive NLuc (x2, x3, x4), and Vpr-fused HiBiT were infected with VSV-G-pseudotyped-NL4-3/KFS. The NLuc activity in the cell and the cell supernatant was measured three days post-infection. The virus and the virus-free cells were fractionated from the cell supernatant. (B) Jurkat cells were infected with VSV-G-pseudotyped-NL4-3/KFS. The amount of p24 was measured three days post-infection. (C) 293T cells encoding the LTR-driven Vpr-fused HiBiT were transfected with pNL4-3/KFS. The NLuc activity in the cell and the cell supernatant was measured two days post-transfection. (D) 293T cells were transfected with pNL4-3/KFS, and the amounts of p24 were measured two days post-transfection. (E) Jurkat cells encoding the LTR-driven indicated luciferase (firefly: FLuc, renilla: RLuc) were infected with VSV-G-pseudotyped- NL4-3/KFS. The luciferase activities were measured three days post-infection. (F) Jurkat/Vpr-HiBiT cells were infected with VSV-G-pseudotyped-NL4-3/KFS. The infected cells were treated with drugs (efavirenz, EFV; nevirapine, NVP) at 0 h and 15 h post-infection. The NLuc activities were measured three days post-infection. The error bars denote the means ± standard deviation of three independent experiments(n=3). The One-way ANOVA by Sidak’s multiple comparison test (B, C, and D) and the Two-way ANOVA using Tukey’s multiple comparison test (A, E, and F) were applied to compare the data. *, *P* < 0.01; **, p<0.001; ***, p<0.0001; *n.s.*, not significant.

### Compounds selected from the library reduce the virus release

To identify the leader compounds that would inhibit virus release, Jurkat/Vpr-HiBiT cells were infected with VSV-G-pseudotyped NL4-3 using the core library from drug discovery initiative (9600 compounds, 5 µM) and the Ono Pharmacy compound library (3280 compounds, 5 µM). The infected cells were treated with these compounds at 15 h post-infection to focus on the late-stage inhibition of viral replication. None of the 9600 compounds in the core library reduced both Vpr-HiBiT and p24 Gag in the cell supernatant (Data not shown). Ninety-four of the 3280 compounds reduced Vpr-HiBiT in the cell supernatant, but 10 out of these 94 compounds did not reduce p24 Gag (Fig. 2A). These results suggest that these 10 compounds would inhibit Vpr assembly into the virions or the binding between HiBiT and LgBiT as described in the discussion. Fifty out of the 94 compounds induced obvious cell toxicity as observed by microscopy. Therefore, the remaining 34 compounds were selected from the first drug screening (red dots in Fig. 2A). In the second drug screening, 10 out of the remaining 16 reduced the nanoluciferase activity without cell toxicity (red dots in Fig. 2B). Seven out of 10 selected compounds consistently reduced the amount of p24 Gag by more than 40% (red dots in Fig. 2C). The compound ONO#05 markedly reduced the p24 Gag in the cells and the cell supernatant (Fig. 2D), a result suggesting that ONO#05 would either inhibit Gag protein synthesis or induce the death of HIV-1-infected cells through the lock-in and apoptosis pathway [51]. Nonetheless, the compound ONO#08 reduced p24 Gag to about 60% in the cell supernatant without inhibiting the amount of p24 Gag in the cells.

**Fig. 2.**
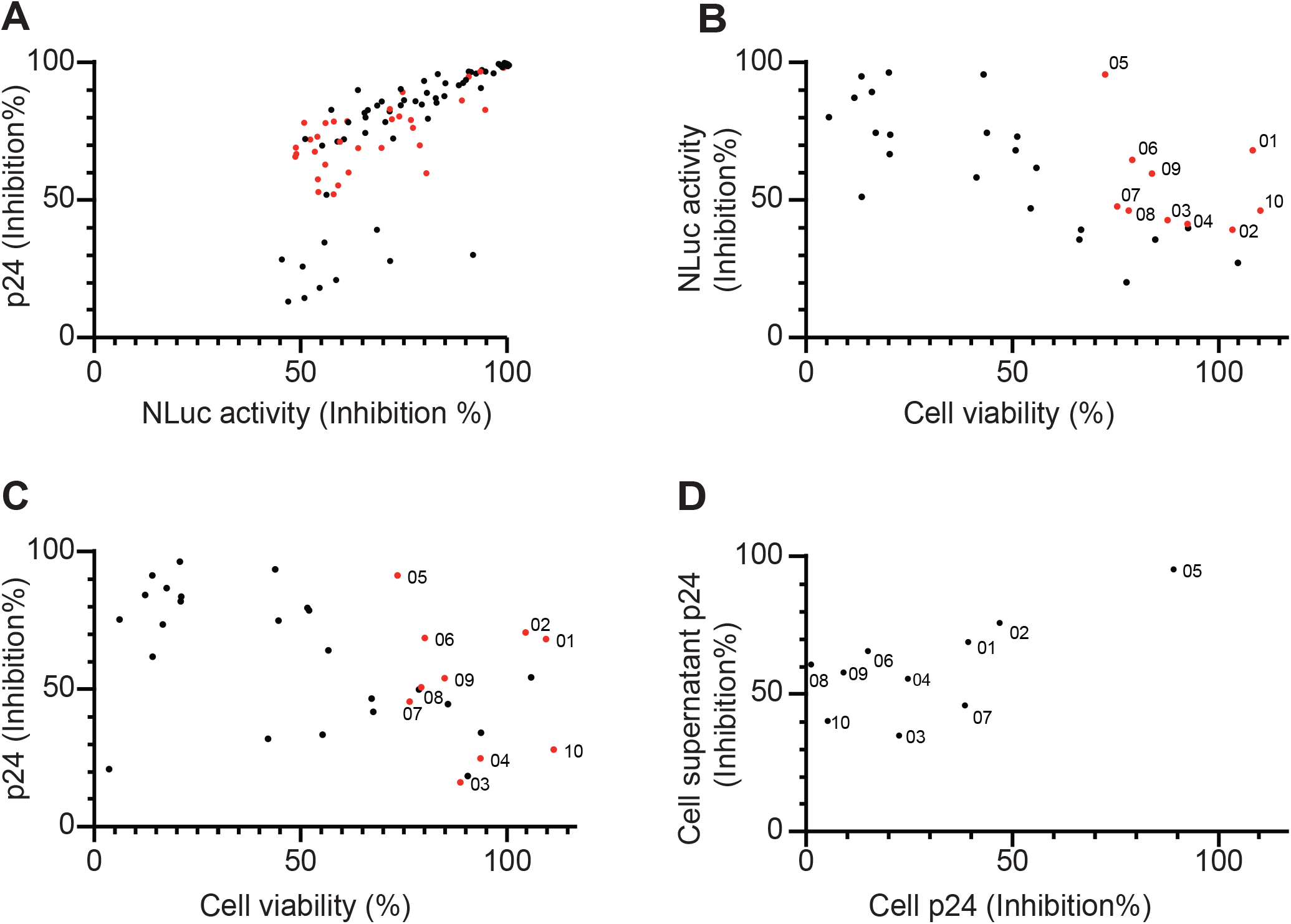
Ten small compounds are candidates for virus release inhibition. (A - D) Jurkat/Vpr-HiBiT cells were infected with VSV-G-pseudotyped-NL4-3/KFS. The infected cells were treated with the compounds selected from the Ono Pharmaceutical compound library (drug discovery initiative of the Tokyo University) at 15 h post-infection. The NLuc activity and the amount of p24 in the cell supernatants (A - D) and the cells (D) were measured three days post-infection. (B and C) The uninfected cells were treated with the compounds. Two days after compound treatment, the cell viability was examined by MTT assay.

### Derivative compound HT-7 reduces virus release

Three compounds (ONO#06, #07, #08), out of the ten selected, had a benzothiophene dioxide (BTD group), as indicated by the red dotted square (Fig. 3A); we focused on ONO#08 to design the derivatives (Supplementary Fig. S1). Initially, the region of disparate structures among the selected compounds (ONO#06, #07, #08) was modified to form derivative compounds (Fig. 3B). However, these derivatives exhibited high cellular toxicity (Fig. 3D). As a result, the derivative compounds (HT-5, HT-6, and HT-7) were designed by changing the BTD group on the leader compound (ONO#08) (Fig. 3C and Supplementary Fig. S1). Unexpectedly, HT-1, which has the same structure as ONO#08, reduced Vpr-HiBiT in the cell supernatant only by 25% and caused cytotoxicity (Fig. 3E). It suggests that the purity of ONO#08 from Ono Pharmacy compound library might have decreased due to its prolonged storage. Importantly, the derivative HT-7 (50 µM) reduced Vpr-HiBiT in the cell supernatant by 60% and caused lower cytotoxicity (Fig. 3E). The EC_50_ and CC_50_ values for HT-1 were 1.4 µM and 4.4 µM, respectively (Fig. 3F). Compared to those for HT-1, the EC_50_ and CC_50_ values for HT-7 were 95.8 µM and 418.9 µM, respectively (Fig. 3F). The leader compound (ONO#08) did not reduce the expression of HIV-1 Gag in the cells (Fig. 2D), suggesting that these derivatives (HT-5, HT-6, and HT-7) would inhibit viral release from HIV-1-infected T cells.

**Fig. 3.**
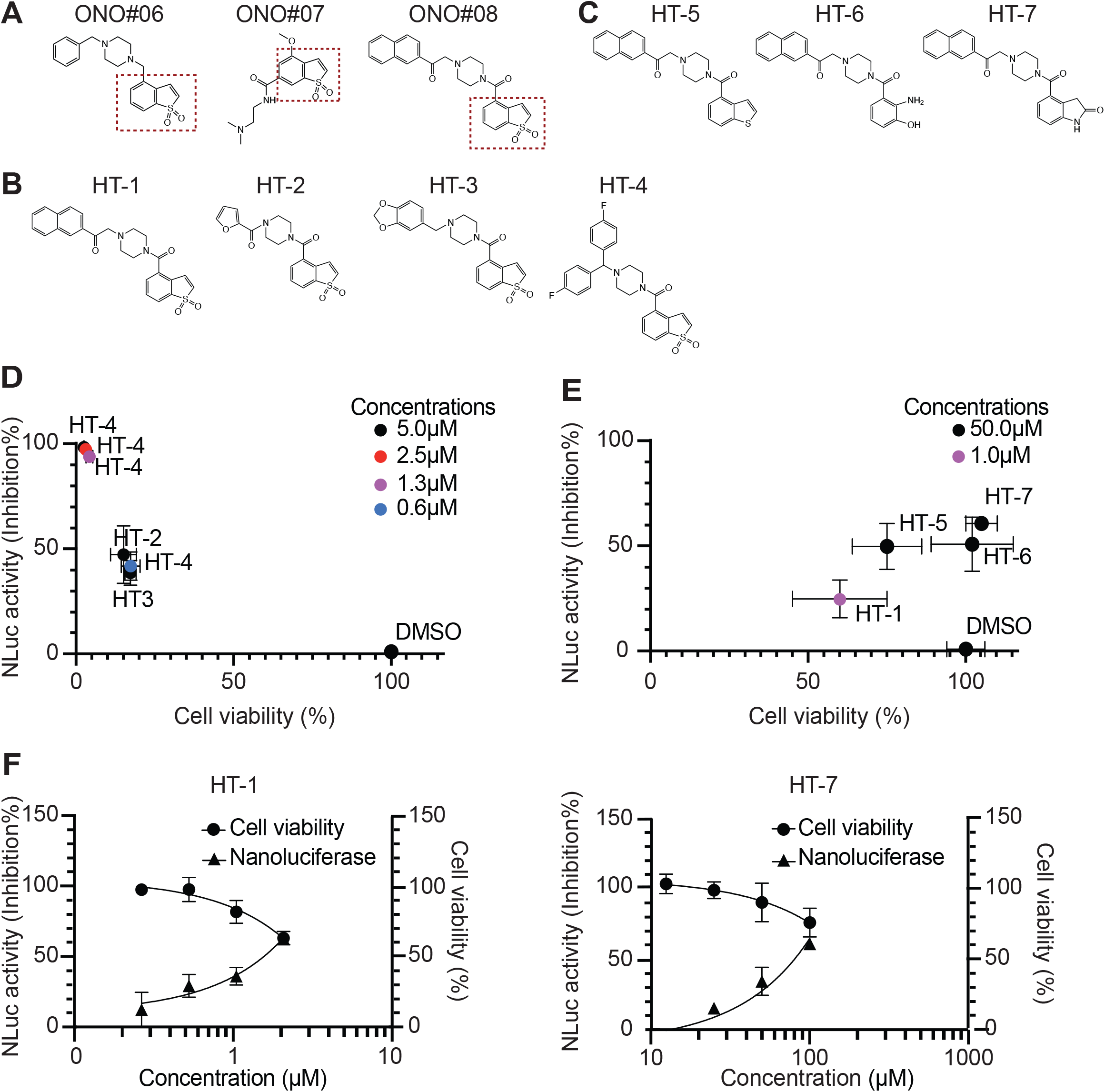
Derivatives inhibit the virus release. (A - C) Structures of the three candidates and nine derivative compounds. A BTD group is shown in the purple dot box. (D and E) Jurkat/Vpr-HiBiT cells were infected with VSV-G-pseudotyped-NL4-3/KFS. The infected cells were treated with the indicated compounds at 15 h post-infection. The NLuc activity was measured three days post-infection. The uninfected cells were treated with the indicated compounds. Two days after compound treatment, the cell viability was examined by MTT assay. (F) The Prism software determined EC_50_ and CC_50_.

### Gag accumulation in the sites of cell polarity is disturbed by HT-7

Following the binding of HIV-1 Gag to the plasma membrane (PM), it is assembled at the PM. To assess the impact of HT-7 on HIV-1 Gag expression and localization to the PM, Jurkat/Vpr-HiBiT cells were infected with VSV-G-pseudotyped NL4-3-GagVenus. HT-7 significantly enhanced the mean fluorescence intensity (MFI) of GagVenus in the cells compared to that in DMSO-treated cells (DMSO; MFI=474, HT-7; MFI=641) (Figs. 4A and 4B). These results suggest that the derivative compounds might induce Gag accumulation into the cells or accelerate Gag expression in the cells. Gag localization at the PM was approximately 40% in Jurkat/Vpr-HiBiT cells with DMSO (Figs. 4C and 4D). Moreover, confocal microscopy showed approximately 20% cell polarity in DMSO condition, observed by Gag accumulation at the uropod surface (Figs. 4C and 4E). Compared to DMSO, HT-7 slightly increased the Gag localization at the PM (to approximately 60%) (Fig. 4D).

**Fig. 4.**
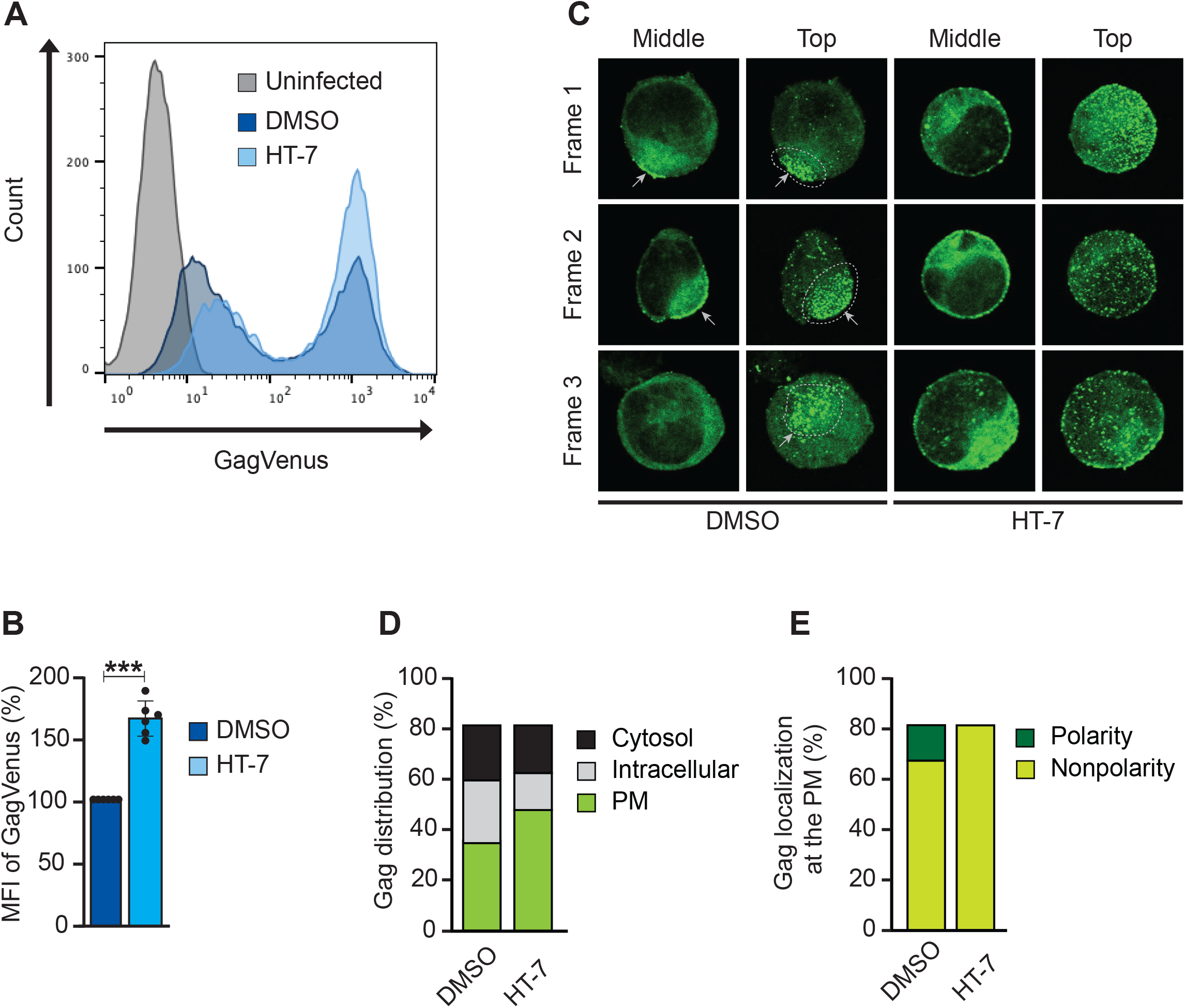
Gag localization is affected by HT-7. Jurkat/Vpr-HiBiT cells were infected with VSV-G-pseudotyped-NL4-3/GagVenus. The infected cells were treated with 50 µM HT-7 at 15 h post-infection. (A - B) The cells were analyzed by Flow cytometry. (C) The cells were observed using confocal microscopy. Images were acquired at the mid-section and the top-section of the cells. Gray arrows indicate Gag accumulation in the polarized area. (D) The cells with Gag localized predominantly to the PM (green), to the intracellular compartments (gray), or in the cytosol (black) were counted. About 100 cells that showed Gag signals were examined. (E) The polarized (dark green) or nonpolarized (light green) cells with Gag localized to the PM were counted. Each cell was imaged from top to bottom and the number of cells with Gag accumulated on the plasma membrane was counted.

Nevertheless, HT-7 perturbed the cell polarity, resulting in Gag distribution all over the cell surface (Fig. 4C and 4E). These results suggest that HT-7 induces Gag accumulation in infected cells by changing the cell membrane state, such as membrane fluidity and/or cytoskeleton-like actin filaments.

### HT-7 does not regulate the cell surface expression of PSGL-1

We postulated that HT-7 might influence the expression and localization of cell surface microdomains, such as P-selectin glycoprotein ligand-1 (PSGL-1), CD43, and CD44, due to its effects on the membrane state. However, the expression of PSGL-1 on the cell surface was not regulated by HT-7 treatment (Fig. 5A). Previous studies have proved that HIV-1 Gag co-assembles with PSGL-1 via the highly basic region (HBR) of MA [52]. In pursuit of disclosing HT-7’s implications in HIV-1 Gag engagement with PSGL-1, we adopted HIV-1 constructs designated as Fyn(10), which has two palmitoyl groups for stable membrane binding. The Fyn(10)/6A2T mutant lacks a HBR in the MA domain and binds to the plasma membrane regardless of PI(4,5)P_2_. Regardless of using these mutants, HT-7 did not affect the co-assembly between HIV-1 Gag and PSGL-1 (Figs. 5B and 5C). Akin to the findings from a previous report [52], we detected an inverse relationship between PSGL-1 expression and HIV-1 Gag expression and a direct association between PSGL-1 expression and Fyn(10)/6A2T Gag expression (Fig. 5D). However, HT-7 did not appear to have any effect on the correlation between HIV-1 Gag and PSGL-1’s in Fyn(10)/6A2T (Fig. 5D).

**Fig. 5.**
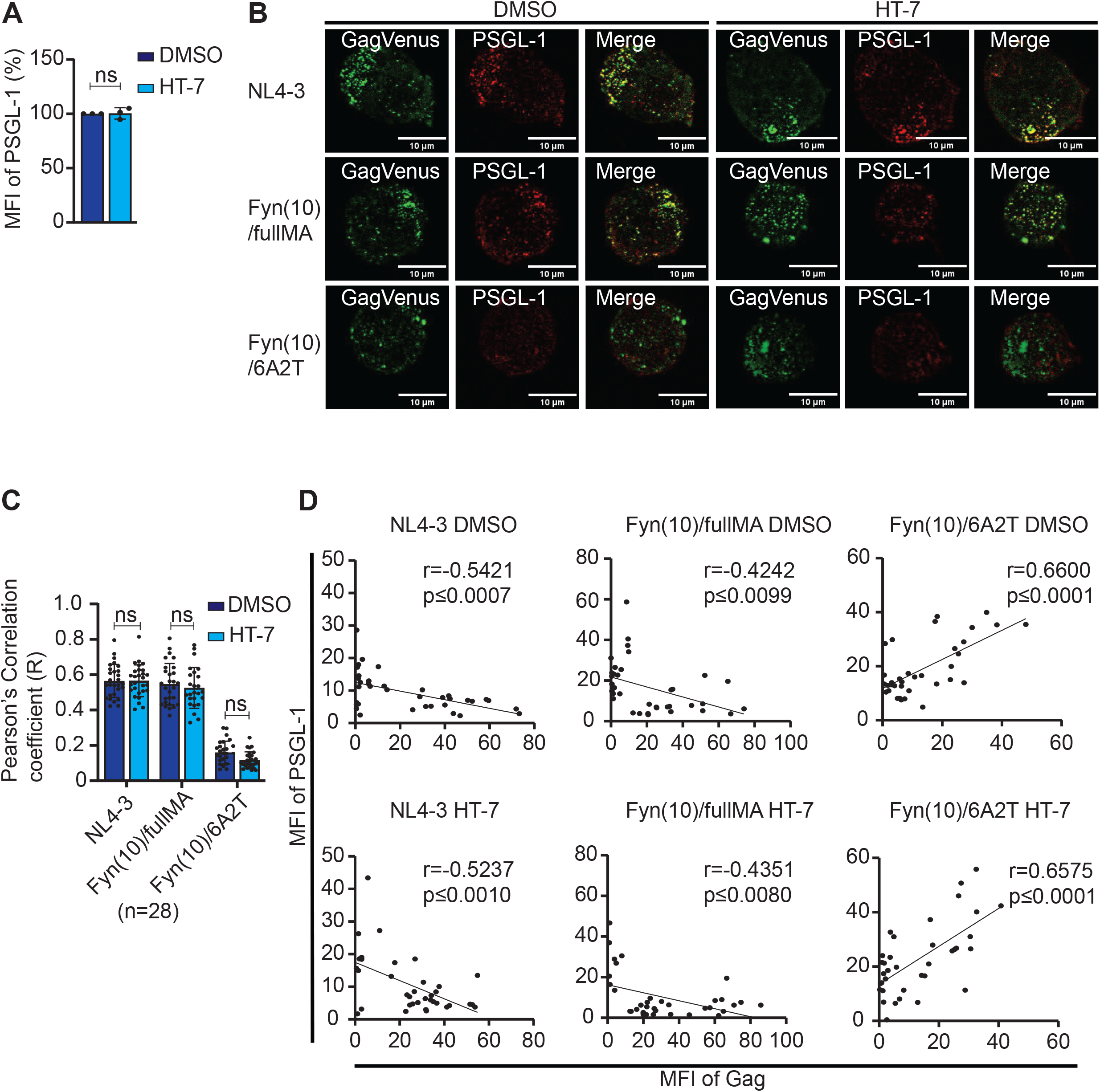
Cellular surface expression of PSGL-1 nor its interaction with HIV-1 Gag is not affected by HT-7. (A) Jurkat cells were infected with VSV-G-pseudotyped-NL4-3/GagVenus for 15 h, after which infected cells were washed and then treated with 50 µM HT-7. The infected cells were incubated for 48 h, then cells were analyzed by flow cytometry. The error bars denote the means ± standard deviation of three independent experiments(n=3). The One- way ANOVA by Sidak’s multiple comparison test (B, C, and D) and the Two-way ANOVA using Tukey’s multiple comparison test (A, E, and F) were applied to compare the data. *, *P* < 0.01; **, p<0.001; ***, p<0.0001; *n.s.*, not significant. (B) Representative diagram of HIV- 1 Gag co-clustering with PSGL-1 in the presence or absence of HT-7. Jurkat/Vpr-HiBiT cells were infected with or without VSV-G-pseudotyped-NL4-3/GagVenus, VSV-G-pseudotyped- Fyn(10)/fullMA/GagVenus and VSV-G-pseudotyped-Fyn(10)/6A2T/GagVenus for 15 h, and the cells were observed by confocal laser microscopy. The green color indicates Gag-YFP signals and the red color indicates PSGL-1 signals. (C) Each dot denotes single cells, n=28. Error bars indicate the mean ± standard deviation. The correlation (r) between the MFI of PSGL-1 and HIV-1 Gag in control and treated conditions was calculated by Pearson’s correlation test. *P* values were determined by Two-way ANOVA using Tukey’s multiple comparison test. *, *p* < 0.01; **, *p* < 0.001; ***, *p* < 0.0001; *n.s.*, not significant. (D) By employing Image J software, the MFI of HIV-1 GagVenus and PSGL-1 was determined. Each dot denotes an individual cell, n=36. Error bars show the means ± standard deviation of repeated tests. The correlation (r) between the MFI of PSGL-1 and HIV-1 Gag in control and treated conditions was calculated by Pearson’s correlation test. *, *p* < 0.01; **, *p* < 0.001; ***, *p* < 0.0001; *n.s.*, not significant.

### Cell surface expression of BST2 is upregulated upon HT-7 treatment

Because of these findings, we re-directed our assumption to other plasma membrane-associated molecules. BST2/Tetherin/CD317/HM1.24 is a well-known restriction factor that inhibits HIV-1 release by tethering the virus onto the plasma membrane. Besides, HIV-1 Vpu counteracts the anti-release effect of BST2 [31]. Given the above, we analyzed BST2 cell surface expression in Jurkat/Vpr-HiBiT cells treated with HT-7. The treatment of uninfected Jurkat/Vpr-HiBiT cells and NL4-3-infected Jurkat/Vpr-HiBiT cells with HT-7 led to a significant increase in MFI of BST2 on the cell surface (Figs. 6A and 6B). Contrarily, in Vpu-deficient NL4-3, BST2 expression on the cell surface was not significantly increased by HT-7 treatment (Fig. 6B). Consequent to the treatment of HT-7 to cells infected with HIV-1 mutants lacking Vpu, no further decline in NLuc activity and p24 amounts was determined (Figs. 6C and 6D). These results suggest that HT-7 reduces the viral release by canceling the effects of Vpu, thereby retaining BST2 on the cell surface.

**Fig. 6.**
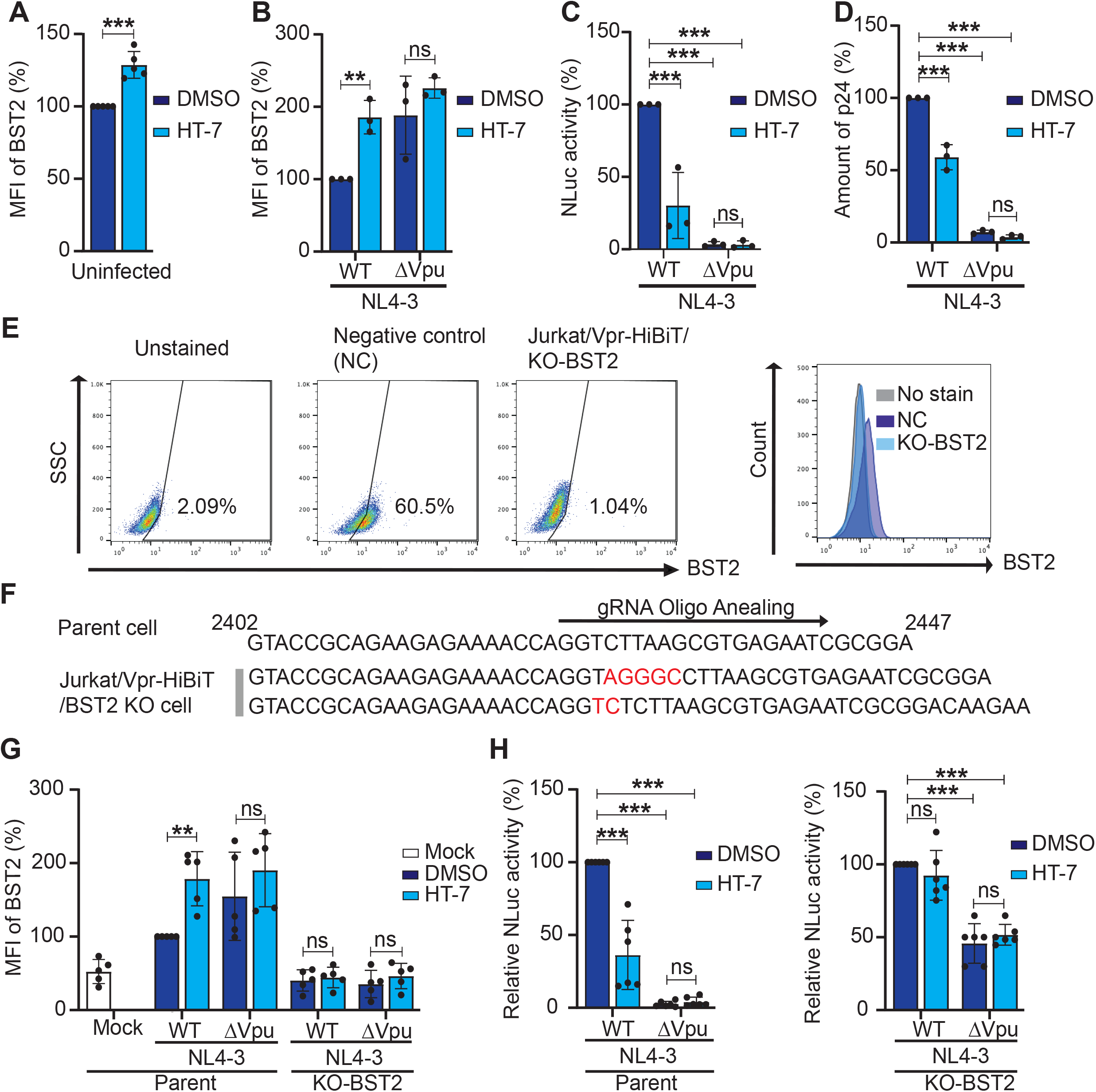
Knockout of BST2 canceled HT-7’s release inhibitory effect. The MFI of BST2 on Jurkat/Vpr-HiBiT cells (A) and infected Jurkat/Vpr-HiBiT cells (B) was determined following analysis by flow cytometry. The NLuc activity (C) and the amount of p24 (D) in the supernatant from the infected Jurkat/Vpr-HiBiT cells were analyzed three days post- infection. Error bars show the mean ± standard deviation of repeated independent experiments. (E) The BST2-positive cell population was analyzed by flow cytometry. (F) The sequences of BST2 in Jurkat/Vpr-HiBiT and Jurkat/Vpr-HiBiT/KO-BST2 cells were analyzed by Sanger sequencing. A guide RNA for BST2 knockout is designed as shown by the arrow. The inserted nucleotides are shown as red color. (G) The MFI of BST2 on Jurkat/Vpr-HiBiT or Jurkat/Vpr-HiBiT/BST2-KO cells was determined following analysis by flow cytometry. (H) The NLuc activity in the supernatant from the infected Jurkat/Vpr-HiBiT or Jurkat/Vpr-HiBiT/KO-BST2 cells was analyzed three days post-infection. (C, D, G, and H) The error bar illustrates the mean ± standard deviation of the repeated independent experiment. Direct comparisons between control and treated conditions were done by Two-way ANOVA using Tukey’s multiple comparison test. *, *p* < 0.01; **, *p* < 0.001; ***, *p* < 0.0001; *n.s.*, not significant.

### BST2 knockout abolished HT-7’s suppressive effect on virus release

To confirm whether the effect of virus release suppression is due to the induction of BST2 by HT-7 treatment, we established knockout BST2 cells using the CRISPR/Cas9 mediated genome editing technique (Figs. 6E and 6F). In Jurkat Vpr-HiBiT BST2 knockout cell lines (Jurkat/Vpr-HiBiT/KO-BST2), five (5) and two (2) nucleotides were inserted into the open reading frame of the *bst2* gene. HT-7 treatment did not upregulate BST2 expression in Jurkat/Vpr-HiBiT/KO-BST2 cells (Fig. 6G). Furthermore, the inhibitory effect of HT-7 was abolished in the absence of BST2 (Fig. 6H). In addition, HT-7 did not further reduce the Vpu-defective mutant in the BST2 knockout cells (Fig. 6H). Notably, we discovered a significant decline of about 50% in the release of Vpu defective mutants even in the BST2 knockout cells (Fig. 6H). These results suggest the possibility of another yet-to-be-discovered role of Vpu in viral replication. Thus, there is the possibility of another cellular restriction factor whose activities may be hindered by Vpu. Otherwise, Vpu might recruit the cellular cofactor for the virus release. These data demonstrate that the attenuation of HIV-1 release by HT-7 is the sequel to its enhancement of BST2 expression on the surface of T-cell lines.

### HT-7 alleviates SIVmac239 release and SARS-CoV-2 replication

Earlier studies have extensively unveiled the obstructive influence of BST2 release on other retroviruses, such as simian immunodeficiency virus (SIV), likewise distinct viral counteractive measures directed against BST2. Concomitantly, we hypothesized that HT-7 interferes with viral release in SIVmac239. To analyze SIVmac239 release from infected T cells, we measured the amounts of p27 Gag in virions (Fig. 7A). We found that HT-7 reduced p27 Gag (Fig. 7A), suggesting that HT-7 broadly inhibits the release of numerous viruses by induction of BST2 on the cell surface.

**Fig. 7.**
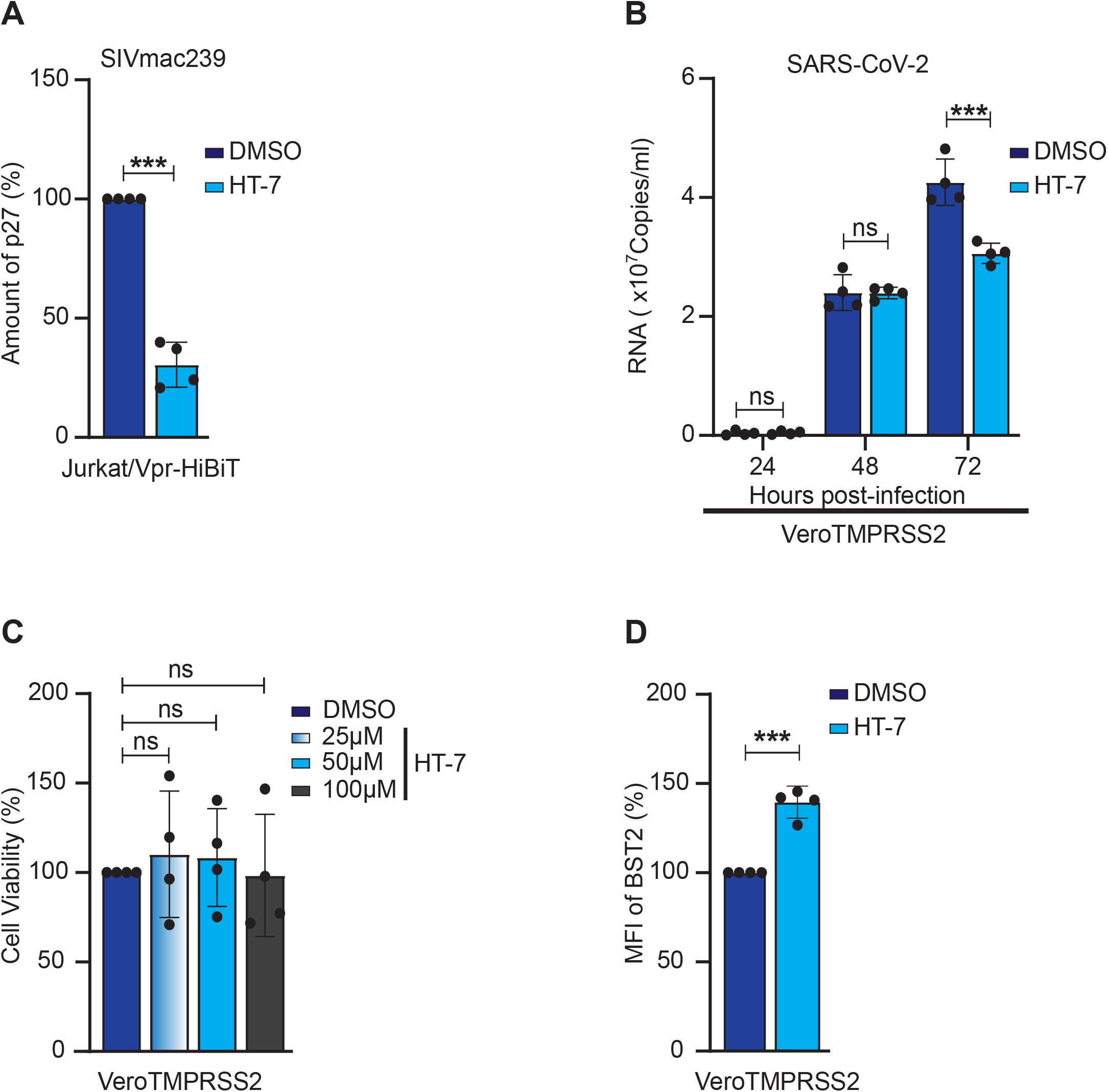
HT-7 alleviates SIVmac239 and SARS-CoV-2 release. (A) Jurkat Vpr-HiBiT cells were infected with VSV-G-pseudotyped SIVmac239ΔEnv for 15 h. Infected cells were treated with 50 µM HT-7. After two days of incubation, p27 amounts in the cell supernatant were analyzed. (B) VeroE6/TMPRSS2 cells were infected with SARS-CoV-2 B.1.1 and were treated with 50 µM of HT-7. At 0, 24, 48, and 72 hrs post-infection, RNA copy in the cell supernatant was quantified by RT-qPCR. (C) The uninfected VeroE6 TMPRSS2 cells were treated with HT-7 (25 µM, 50 µM, and 100 µM). Three days after HT-7 treatment, the cell viability was examined by MTT assay. (D) The MFI of VeroE6 TMPRSS2 cells was analyzed by flow cytometry. Error bars show the mean ± standard deviations of four experiments. Comparisons of control and treated conditions were evaluated by the unpaired *t*- test (A and D), Two-way ANOVA using Tukey’s multiple comparison test (B), and One-way ANOVA using Sidak’s multiple comparison test (C). *, *p* < 0.01; **, *p* < 0.001; ***, *p* < 0.0001; *n.s.*, not significant.

In our quest to ascertain the efficacy of HT-7 against other enveloped viruses, we assessed its impacts on SAR-CoV-2 replication. SARS-CoV-2 harnesses its ORF3a and spike protein to counteract the effect of BST2 on its replication [53, 54]. We detected a significant reduction in SARS-CoV-2 replication by HT-7 treatment at 72 h post-infection without cell toxicity (Fig. 7B and 7C). Furthermore, we confirmed the increment of cellular surface BST2 by HT-7 in VeroE6/TMPRSS2 cells (Fig. 7D). These results signify that similar to HIV-1, HT-7 reduces SARS-CoV-2 release by augmenting BST2 on the cell surface.

## Discussion

In this study, a novel high-performance technique (Vpr-HiBiT screening) for monitoring the efficiency of budding virions in T-cell lines was established. Using this Vpr-HiBiT screening system, we discovered 10 leader compounds from the Ono Pharmaceutical compound library. Here, we revealed that the derivative compound, HT-7, designed from selected candidate compounds, inhibited HIV-1 release in a T cell line as well as SIVmac239 release and SARS-CoV-2 replication by increasing cell surface BST2. Further, in the presence of HT-7, HIV-1 accumulated in producer cells.

In this screening system, NLuc activity based on the Vpr-HiBiT indirectly reflects the amounts of virions in the supernatant from T cell lines. However, in 293T/Vpr-HiBiT cells, Vpr-HiBiT did not reflect the amounts of virions because half of the Vpr-HiBiT was virus-free in the cell supernatant. The mechanism of cell-line-specific Vpr-assembly is not yet known precisely. Nonetheless, the small peptide HiBiT is crucial for the efficient assembly of Vpr into virions. The HiBiT peptide needs to fit into the binding pocket of LgBiT for detection of NLuc activity [55]; therefore, compounds, such as DrkBiT [56], interfering with the binding between HiBiT and LgBiT may be selected as false candidates. Hence, to avoid the selection of incorrect candidates, direct quantification, which measures p24 Gag in the cell supernatant, is essential for determining candidate compounds by secondary screening.

Unexpectedly, we were unable to discover any Gag inhibitors in this study. The reason may be that since Gag is expressed in cells in large quantities and is assembled into virions [57, 58], a large amount of small compounds may be necessary for blocking the Gag-Gag interaction in cells. Therefore, identifying small compounds that interfere with Gag multimerization at the PM might be challenging. It suggests that compounds that target the host molecules might be selected more predominantly through this screening system.

Currently, lenacapavir, a compound that fits into the binding pocket of the Gag CA domain, is in the spotlight as an inhibitor of the post-entry step [6, 28]. The structure of lenacapavir is complex and completely different from that of the small compounds in the general drug library. It suggests that we need to search for candidates from natural compounds to target the Gag protein. Alternatively, in silico molecular docking studies might be an alternative approach to identifying complex compounds.

The derivative compound, HT-7, with considerable release inhibition and inappreciable cytotoxicity, remarkably elevates the surface expression of BST2 but not PSGL-1. Since BST2 co-assembles with viral structural proteins at the viral forming regions of the PM and tethers the virions on the cell surface, it is likely that HT-7 broadly impedes viral release in enveloped viruses. Predictably, HT-7 attenuation of SIVmac239 release and SARS-CoV-2 replication emphasizes its broadly restraining effect against the release of other enveloped viruses. Whether the influence of HT-7 on BST2 surface expression involves indirect or direct contact with Vpu is unclear. Notwithstanding, viral restriction factors against BST2 differ among enveloped varies. Hence, the upregulation of BST2 expression by HT-7 may not be associated with a direct interaction of HT-7 with viral counteractive proteins.

As a future direction, whether HT-7 binds directly to BST2 should be investigated. Studies in the past proved that BST2 can ensnare exosomes into plasma membranes just as enveloped viruses [59]. The potency of intact BST2 but not a BST2 deficitient in GPI anchor to influence extracellular exosome levels has also been reported [59, 60]. Accordingly, it suggests that a novel therapeutic agent designed from HT-7 may be able to cure several disorders related to exosomes via its augmentation of cell surface BST2. For example, the release of exosomes, which are associated with neurological diseases and the progression of malignant diseases [61, 62], might also be suppressed by HT-7. Furthermore, BST2 is one of the markers of cancer cells and is a target for anticancer therapy by the induction of antibody-dependent cellular cytotoxicity. Thus, HT-7 may be expected to serve as a broad-range therapeutic agent.

Ultimately, the established high-performance screening technique, the Vpr-HiBiT system, can be employed in screening late-stage inhibitors. Moreover, it is economical and requires an unusually short processing and testing time, emphasizing its usefulness for basic research. Further studies to design alternative derivative compounds with effective inhibition ability in primary cell lines and animal models need to be undertaken.

## Acknowledgments

This research was supported by the Platform Project for Supporting Drug Discovery and Life Science Research from AMED under Grant Number JP20am0101086 (support number 1905). The authors thank Ono Pharmaceutical Co., Ltd. and the Tokyo University Drug Discovery Initiative for providing these compounds. This work was supported by AMED (21fk0410026h0001, 24fk0410065h0001) to K.M., JSPS KAKENHI (19K23802) to H.T., AMED (22fk0410052h0001) to Y.S., and Young Investigator Award at Joint Research Center for Human Retrovirus Infection, Kumamoto University to K.M. We would like to thank Editage (www.editage.com) for English language editing, and thank Dr. E. Freed for providing plasmids. The following reagent was obtained through the AIDS Research and Reference Reagent Program, Division of AIDS, NIAID, NIH: SIVmac239 infectious Molecular Clone (SIVmac239 M5 SpX), ARP-12247, contributed by Dr. Ronald C. Desrosiers.

## Author’s Contributions

KM, HT,YM, MF, YS, TI, TS, and MO conceived and coordinated the study. PN, AT, TY, TM, MMB, WS, KS, HT, NM, YT, JAK, WOA, MJH, and KM performed the experiments.

KM and TY established the cell lines for drug screening. RT and YT established the p24 ELISA system. KY isolated the SARS-CoV-2. HT and NM supported the drug screening.

AK and HT developed the derivative compounds. All the authors have read and approved the final manuscript.

## Declaration of Interests

The authors declare that they have no competing interests.

---

**Supplemental Fig. S1.**
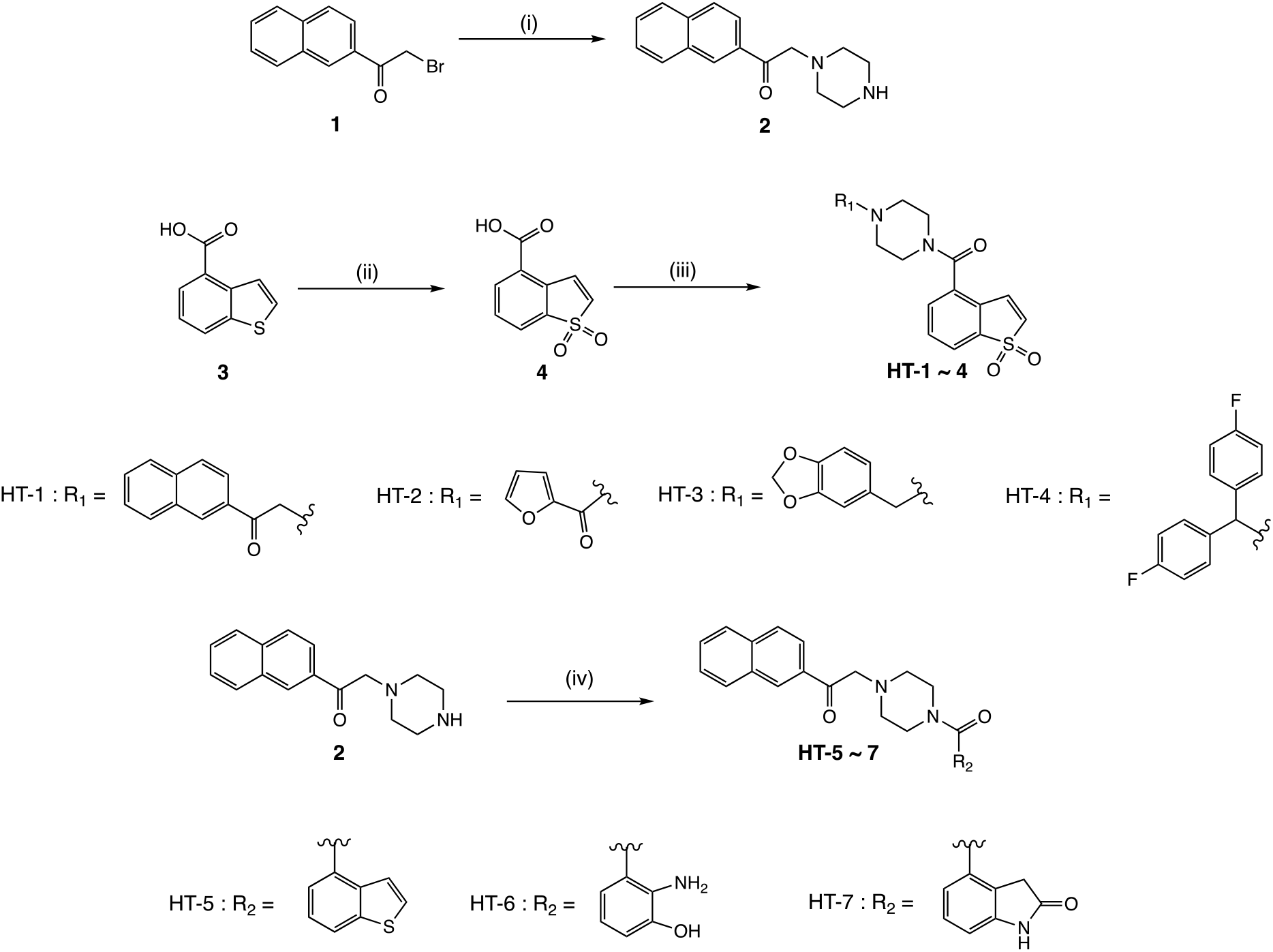
Scheme 1. Synthesis of HT 1 ∼ 7 Reagents and conditions: (i) piperazine, EtOH, reflux, overnight, (ii) oxone, MeOH, 40 ℃, 2 h, (iii) a) oxalyl chloride, CH_2_Cl_2_, reflux, 3 h, b) **2**, Et_3_N, CH_2_Cl_2_, r.t., 2 h, (iv) DMT-MM, MeOH, r.t., 3 h.y

## Experimental

### General procedure for the synthesis

All reagents were commercially available and of high-grade purity. Thin layer chromatography (TLC) was performed on precoated plates (TLC sheets silica 60 F254) (Merck, Darmstadt, Germany). Chromatography was performed on Silica Gel 60N (40–100 mesh) (Kanto Chemical, Tokyo, Japan). NMR spectra were recorded on a Bruker AVANCE 600 (Billerica, MA, USA). The chemical shifts were referenced to TMS. Mass spectrum fast atom bombardment (FAB) and high-resolution mass spectra (HRMS) were recorded by a JEOL JMS-DX303 (Tokyo, Japan). HRMS were recorded by using positive FAB with 3- nitrobenzyl alcohol (NBA) as the matrix.

### Synthesis

1-(naphthalen-2-yl)-2-(piperazin-1-yl) ethan-1-one (**2**):

Bromide compound **1** (3.9 g, 16 mmol) was dissolved in 50 mL EtOH, 2- piperazine (6.7 g, 78 mmol) and DIPEA (2.2 ml, 16 mmol) was added, and the mixture was refluxed overnight. The solvent was removed and the residue was dissolved in AcOEt (50 mL) and washed with H_2_O (50 mL × 2). The organic layer was concentrated under reduced pressure and purified by column chromatography (AcOEt: MeOH: Et_3_N = 1:10:0.1). **2** (2.0 g, 60%) was obtained as a brown oil [1]. ^1^H NMR (600 MHz, CDCl_3_) δ (ppm): 2.64 (4H, bs, CH_2_ x 2), 2.85 (1H, bs, NH), 2.99 (4H, bs, CH_2_ x 2), 3.92 (2H, s, CH_2_), 7.54 (1H, t, *J* = 7.2 Hz, CH), 7.59 (4H, t, *J* = 7.0 Hz, CH), 7.87 (2H, t, *J* = 9.0 Hz, CH x 2), 7.95 (1H, d, *J* = 8.0 Hz, CH), 8.04 (1H, d, *J* = 8.4 Hz, CH), 8.55 (1H, s, CH). ^13^C NMR (150 MHz, CDCl_3_) δ 45.8, 54.6, 65.2, 123.9, 126.8, 127.8, 128.4, 128.5, 129.6, 129.8, 132.5, 133.5, 135.7, 196.4. HRMS (FAB) m/z calcd for C16H19N2O (M + H)^+^ 255.1497. Found: 255.1497. TLC: Rf 0.30 (AcOEt:MeOH: Et_3_N = 1:10:0.1).

benzo[b]thiophene-4-carboxylic acid 1,1-dioxide (**4**):

**3** (0.16 g, 0.90 mmol) was dissolved in MeOH (10 mL), Oxone (1 g, 6.6 mmol) in water (10 mL) was added, and stirred at 40 ℃ for 2 h. The reaction mixture was extracted with EtOAc (20 mL), dried over Na_2_SO_4_, and concentrated under reduced pressure. The residue was suspended in hexane and collected precipitation **4** (160 mg, 85%) as a white solid (1).

^1^H NMR (600 MHz, MeOD) δ 7.12 (1H, d, *J* = 7.1 Hz, CH), 7.68 (1H, t, *J* = 7.7 Hz, CH), 7.91 (1H, d, *J* = 7.6 Hz, CH), 8.27 (1H, dd, *J* = 1.0, 7.9 Hz, CH), 8.28 (1H, dd, *J* = 0.6, 7.2 Hz, CH). ^13^C NMR (150 MHz, MeOD) δ 125.5, 128.4, 131.9, 133.0, 133.1, 133.2, 136.4, 139.3, 167.5. HRMS (FAB) m/z calcd for C_9_H_5_O_4_S (M - H)^-^ 208.9909. Found: 208.9913. TLC: Rf 0.18 (AcOEt).

2-[4-(1,1-dioxidobenzo[b]thiophene-4-carbonyl) piperazin-1-yl]-1-(naphthalen-2-yl) ethan-1-one (**HT-1**)

To the solution of **4** (0.068 g, 0.32 mmol) in CH_2_Cl_2_ (5 mL) was added oxalyl chloride (0.056 mL, 0.65 mmol) and DMF (3 drops), and the mixture was refluxed for 3 h. The solvent was removed and the residue was dissolved in CH_2_Cl_2_ (5 mL), added **2** (0.082 g, 0.32 mmol) and Et_3_N (0.045 mL, 0.32 mmol), stirred for 2 h. The reaction was concentrated under reduced pressure, and the residue was purified by silica gel column chromatography (AcOEt) to afford **HT-1** (0.045 g, 47%) [1].

^1^H NMR (600 MHz, CDCl_3_) δ 2.64 (2H, bs, CH_2_), 2.82 (2H, bs, CH_2_), 3.45 (2H, bs, CH_2_), 3.94 (4H, bs, CH_2_), 4.05 (2H, s, CH_2_), 6.80 (1H, d, *J* = 7.0 Hz, CH), 7.36 (1H, dd, *J* = 0.7, 6.9 Hz, CH), 7.36 (1H, dd, *J* = 0.9, 7.8 Hz, CH), 7.56-7.59 (2H, m, CH x 2), 7.61-7.64 (1H, m, CH), 7.76 (2H, d, *J* = 7.5 Hz, CH), 7.89 (1H, d, *J* = 8.1 Hz, CH), 7.91 (1H, d, *J* = 8.6 Hz, CH), 8.01 (1H, dd, *J* = 1.7, 8.6 Hz, CH), 8.50 (1H, s, CH). HRMS (FAB) m/z calcd for C_25_H_23_N_2_O_4_S (M + H)^+^ 447.1379. Found: 447.1374. TLC: Rf 0.58 (AcOEt). (1,1-dioxidobenzo[b]thiophen-4-yl) [4-(furan-2-carbonyl) piperazin-1-yl]methanone

(**HT-2**):

**2** (0.070 g, 0.33 mmol) and furan-2-yl(piperazin-1-yl)methanone (0.072 g, 0.33 mmol) were allowed to react under the same conditions as described for the preparation of **HT-1** to give **HT-2** (0.085 g, 69%).

^1^H NMR (600 MHz, CDCl_3_) δ 3.45 (2H, bs, CH_2_), 3.77 (2H, bs, CH_2_), 3.90 (2H, bs, CH_2_), 3.94 (2H, bs, CH_2_), 6.51 (1H, dd, *J* = 1.7, 3.3 Hz, CH), 6.82 (1H, d, *J* = 7.0 Hz, CH), 7.08 (1H, d, *J* = 3.4 Hz, CH), 7.36 (1H, d, *J* = 7.1 Hz, CH), 7.50-7.52 (2H, m, CH x 2), 7.60 (1H, t, *J* = 7.6 Hz, CH), 7.78 (1H, d, *J* = 7.5 Hz, CH). ^13^C NMR (150 MHz, CDCl_3_) δ 42.4, 47.5, 111.6, 117.5, 122.4, 129.5, 130.0, 130.8, 131.3, 131.7, 132.0, 137.7, 144.1, 147.4, 159.1, 166.0. HRMS (FAB) m/z calcd for C_18_H_17_N_2_O_5_S (M + H)^+^ 373.0858. Found: 373.0878. TLC: Rf 0.43 (Hexane:AcOEt = 1:10).

[4-(benzo[d][1,3] dioxol-5-ylmethyl)piperazin-1-yl](1,1-dioxidobenzo[b]thiophen-4- yl)methanone (**HT-3**):

**2** (0.070 g, 0.33 mmol) and 1-(benzo[d][1,3]dioxol-5-ylmethyl)piperazine (0.073 g, 0.33 mmol) were allowed to react under the same conditions as described for the preparation of **HT-1** to give **HT-3** (0.062 g, 45%).^1^H NMR (600 MHz, CDCl_3_) δ 2.35 (2H, bs, CH_2_), 2.52 (2H, bs, CH_2_), 3.32 (2H, bs, CH_2_), 3.44 (2H, s, CH_2_), 3.81 (2H, bs, CH_2_), 5.94 (2H, s, CH_2_), 6.71-6.75 (2H, m, CH x 2), 6.79 (1H, d, *J* = 7.1 Hz, CH), 6.83 (1H, d, *J* = 1.1 Hz, CH), 7.32 (1H, d, *J* = 6.9 Hz, CH), 7.48 (1H, dd, *J* = 0.6, 7.7 Hz, CH), 7.55 (1H, t, *J* = 7.6 Hz, CH), 7.74 (1H, d, *J* = 7.5 Hz, CH). ^13^C NMR (150 MHz, CDCl_3_) δ 42.3, 47.6, 52.5, 53.2, 62.5, 101.0, 108.0, 109.3, 122.0, 122.2, 129.3, 130.2, 130.6, 131.2, 131.3, 131.4, 132.8, 137.5, 146.9, 147.8, 165.6. HRMS (FAB) m/z calcd for C_21_H_21_N_2_O_5_S (M + H) ^+^ 413.1171. Found:413.1169 TLC: Rf 0.50 (Hexane: AcOEt = 1:10).

{4-[bis(4-fluorophenyl) methyl] piperazin-1-yl} (1,1-dioxidobenzo[b]thiophen-4-yl) methanone (**HT-4**):

**2** (0.070 g, 0.33 mmol) and 1-[bis(4-fluorophenyl) methyl] piperazine (0.096 g, 0.33 mmol) were allowed to react under the same conditions as described for the preparation of **HT-1** to give **HT-3** (0.095 g, 59%). ^1^H NMR (600 MHz, CDCl_3_) δ 2.30 (2H, bs, CH_2_), 2.47 (2H, bs, CH_2_), 3.33 (2H, bs, CH_2_), 3.81 (2H, bs, CH_2_), 4.27 (2H, s, CH_2_), 6.77 (1H, d, *J* = 7.0 Hz, CH), 6.96-6.98 (4H, m, CH x 4), 7.30-7.34 (5H, m, CH x 5), 7.45 (1H, dd, *J* = 1.0, 7.8 Hz, CH), 7.51 (1H, t, *J* = 7.6 Hz, CH), 7.70 (1H, d, *J* = 7.5 Hz, CH). ^13^C NMR (150 MHz, CDCl_3_) δ 42.3, 47.7, 51.4, 52.1, 74.1, 115.6, 115.7, 122.0, 129.2, 130.2, 130.6, 131.3, 131.4, 132.6, 137.3, 137.5, 161.1, 162.8, 165.5. HRMS (FAB) m/z calcd for C_26_H_22_N_2_O_3_SNa (M + Na)^+^ 503.1217. Found:503.1211 TLC: Rf 0.50 (Hexane: AcOEt = 1:1). 2-[4-(benzo[b]thiophene-4-carbonyl) piperazin-1-yl]-1-(naphthalen-2-yl) ethan-1- one (**HT-5**):

To the mixture of 2 (0.86 g, 3.4 mmol) and **3** (0.30 g, 1.7 mmol) in MeOH (10 mL) were added DMT-MM (0.56 g, 2.0 mmol), and stirred overnight. After concentrating the reaction mixture under reduced pressure, the residue was dissolved in hexane: AcOEt (1:1, 50 mL) and washed with water. The organic layer was concentrated under reduced pressure, the residue was purified by silica gel column chromatography (Hexane: AcOEt = 1:10) to afford **HT-5** (0.34 g, 49%). After purification, **HT-5** was converted to hydrochloride using a 4M HCl / Dioxane solution.

^1^H NMR (600 MHz, MeOD) δ 3.51-3.80 (8H, m, CH_2_ × 4), 5.21 (2H, s, CH_2_), 7.35 (1H, t, *J* = 7.4 Hz, CH), 7.42-7.43 (2H, m, CH x 2), 7.53 (1H, t, *J* = 7.2 Hz, CH), 7.60 (1H, t, *J* = 7.6 Hz, CH), 7.68 (1H, d, *J* = 5.2 Hz, CH), 7.85-7.96 (4H, CH x 2), 8.00 (1H, d, *J* = 8.0 Hz, CH), 8.59 (1H, s, CH). ^13^C NMR (150 MHz, MeOD) δ 53.9, 55.0, 62.5, 123.1, 124.0, 124.2, 125.2, 125.7, 129.6, 128.5, 129.1, 130.1, 130.4, 130.5, 130.8, 131.0, 132.2, 132.3, 133.8, 137.8, 142.2, 171.4, 191.6. HRMS (FAB) m/z calcd for C_25_H_23_N_2_O_2_S (M + H) ^+^ 415.1480. Found: 415.1492. TLC: Rf 0.47 (Hexane:AcOEt = 1:10).

2-[4-(2-amino-3-hydroxybenzoyl) piperazin-1-yl]-1-(naphthalen-2-yl) ethan-1-one (**HT-6**):

**2** (0.12 g, 0.46 mmol) and 2-amino-3-hydroxybenzoic acid (0.070 g, 0.46 mmol) were allowed to react under the same conditions as described for the preparation of **HT-5** to give **HT-6** (0.064 g, 36%). After purification, **HT-6** was converted to hydrochloride using a 4M HCl / Dioxane solution.

^1^H NMR (600 MHz, MeOD) δ 3.56-3.81 (8H, m, CH_2_ × 4), 5.27 (2H, s, CH_2_), 7.05 (1H, d, *J* = 7.5 Hz, CH), 7.12 (1H, d, *J* = 7.8 Hz, CH), 7.34 (1H, t, *J* = 8.0 Hz, CH), 7.63 (1H, t, *J* = 7.5 Hz, CH), 7.69 (1H, t, *J* = 7.4 Hz, CH), 7.97 (1H, d, *J* = 8.1 Hz, CH), 8.01-8.06 (2H, m, CH x 2), 8.10 (1H, d, *J* = 8.2 Hz, CH) 8.70 (1H, s, CH). ^13^C NMR (150 MHz, MeOD) δ 53.7, 62.6, 119.1, 120.0, 124.0, 128.5, 129.1, 129.9, 130.0, 130.1, 130.8, 131.0, 132.2, 132.3, 133.9, 137.9, 153.0, 168.7, 191.7. HRMS (FAB) m/z calcd for C_23_H_24_N_3_O_3_ (M + H)^+^ 390.1818. Found: 390.1823. TLC: Rf 0.45 (AcOEt MeOH = 19:1). 4-{4-[2-(naphthalen-2-yl)-2-oxoethyl] piperazine-1-carbonyl}indolin-2-one (**HT-7**): **2** (0.14 g, 0.55 mmol) and 2-oxoindoline-4-carboxylic acid (0.097 g, 0.55 mmol) were allowed to react under the same conditions as described for the preparation of **HT-5** to give **HT-7** (0.10 g, 45%). ^1^H NMR (600 MHz, CDCl_3_) δ 2.63 (2H, bs, CH_2_), 2.78 (2H, bs, CH_2_), 3.50 (2H, bs, CH_2_), 3.57 (2H, s, CH_2_), 3.90 (4H, bs, CH_2_), 4.03 (2H, s, CH_2_), 6.92 (1H, d, *J* = 7.9 Hz, CH), 6.95 (1H, d, *J* = 7.4 Hz, CH), 7.24 (1H, t, *J* = 7.7 Hz, CH), 7.55-7.63 (2H, m, CH x 2), 7.88 (1H, d, *J* = 8.2 Hz, CH), 7.90 (1H, d, *J* = 8.6 Hz, CH), 7.96 (1H, d, *J* = 7.9 Hz, CH), 8.02 (1H, dd, *J* = 1.7, 8.6 Hz, CH), 8.51 (1H, s, CH), 8.85 (1H, s, CH). ^13^C NMR (150 MHz, CDCl_3_) δ 54.2, 62.7, 112.6, 124.0, 125.7, 128.1, 128.6, 128.9, 129.1, 129.6, 129.7, 130.2, 130.8, 131.0, 132.2, 132.3, 133.8, 137.9, 145.9, 170.7, 191.6. HRMS (FAB) m/z calcd for C_25_H_23_N_3_O_3_Na (M + H)^+^ 436.1637. Found: 436.1654. TLC: Rf 0.30 (AcOEt MeOH = 19:1).

